# Bayesian Maximum Entropy Ensemble Refinement

**DOI:** 10.1101/2023.09.12.557310

**Authors:** Benjamin Eltzner, Julian Hofstadler, Daniel Rudolf, Michael Habeck, Bert de Groot

**Affiliations:** Max Planck Institute for Multidisciplinary Sciences, Göttingen, Germany; University of Jena, Jena, Germany; University of Passau, Passau, Germany

**Keywords:** Protein Structure, Ensemble Refinement, Maximum Entropy, Adaptive Markov Chain Monte Carlo, Doubly Intractable Problem

## Abstract

The principle of maximum entropy provides a canonical way to include measurement results into a thermodynamic ensemble. Observable features of a thermodynamic system, which are measured as averages over an ensemble are included into the partition function by using Lagrange multipliers. Applying this principle to the system’s energy leads to the well-known exponential form of the Boltzmann probability density. Here, we present a Bayesian approach to the estimation of maximum entropy parameters from nuclear Overhauser effect measurements in order to achieve a refined ensemble in molecular dynamics simulations. To achieve this goal, we leverage advances in the treatment of doubly intractable Bayesian inference problems by adaptive Markov Chain Monte Carlo methods. We illustrate the properties and viability of our method for alanine dipeptide as a simple model system and trp-cage as an example for a more complex peptide.

## 1 Introduction

Observable properties of biomolecules are usually measured on ensembles of many molecules of the same kind. This means that the measured values are ensemble averages and are typically not satisfied by each molecule individually. To take this into account, we present an ensemble refinement method for molecular dynamics (MD) simulations, based on the maximum entropy principle first stated by Jaynes (1957). Following this approach, we achieve a least biased modification of the MD ensemble which satisfies a set of measured ensemble averaged observables. These can in principle be a variety of quantities determined by a variety of measurement methods, usually spectroscopic in nature.

There are numerous ensemble refinement methods based on the maximum entropy principle or related to it, for a review see Cesari et al. (2018). Many approaches focus on *reweighting* the structures in a given prior ensemble, e.g. from an MD trajectory, see Crehuet et al. (2019); Bottaro et al. (2020); Barrett et al. (2022) for some recent examples. Related approaches attempt to achieve better uncertainty treatment using a Bayesian approach, see Hummer and Köfinger (2015); Köfinger et al. (2019), or apply reweighting to entire MD trajectories in a set of multiple trajectories, see Bolhuis et al. (2021). All approaches, based on reweighting a given ensemble, require conformations which are compatible with the measured values of the observable features of interest to already be present in the given ensemble. In most reweighting approaches, while weights for the structures are calculated, no values for the coupling parameters in the maximum entropy ensemble are determined so the probability distribution of the ensemble remains unknown.

An alternative approach to maximum entropy ensemble refinement works directly on the dynamic system by running multiple MD simulations in parallel and applying corrective forces based on the mean values of the observed features across the whole set of simulations. This *replica averaging* approach has been shown to converge to maximum entropy, if the number of replicas goes to infinity, see Pitera and Chodera (2012); Cavalli et al. (2013); Roux and Weare (2013); Boomsma et al. (2014). Since this approach is not dependent on the similarity between a given ensemble and the maximum entropy ensemble, it can be more flexible. However, the convergence to maximum entropy in the limit of infinitely many replicas is theoretical and cannot be practically achieved. Also, replica averaging methods are not capable to provide estimates for the maximum entropy coupling parameters.

A third class of methods aims specifically at estimating the maximum entropy coupling parameters with repeated simulations, see White and Voth (2014); Amirkulova and White (2019). Thereby, these approaches leverage the advantage of replica averaging to not depend on a single given ensemble. However, they are based on a differential treatment of the maximum entropy condition, and are therefore tightly linked to the gradient descent method they employ. Therefore, a Bayesian version of this approach appears to be far from straight forward. Since many MD simulations are performed subsequently, this technique is limited to fairly short MD run time and can therefore be insensitive to conformational dynamics on long time scales.

Estimating the maximum entropy parameters explicitly has several benefits. The maximum entropy parameters can be interpreted as force coupling constants which indicate a modification of the MD force field. Thus, results for different types of measurements can be easily combined by simply adding the corresponding terms to the energy. Since the maximum entropy terms have a natural interpretation as forces, they can be compared to other forces in the system to give a heuristic understanding of the strength of the force field modification. Furthermore, since the measured features concern only parts of a molecule, the maximum entropy parameters include information, which parts of the molecule are modeled well by the MD and at which places the force field is modified. This location information can be used as a starting point for force field refinement beyond the specific system at hand.

In the present paper, we propose an approach for a fully Bayesian treatment of the estimation of maximum entropy coupling parameters. In contrast to other approaches, we tackle the problem that the partition function of the maximum entropy ensemble is not tractable analytically head on. Following the work of Habeck (2014), we perform repeated MD simulations with force fields modified by maximum entropy couplings, like the approach by Amirkulova and White (2019). However, we use the generated MD trajectories to estimate the partition function using the *weighted histogram analysis method* (WHAM) algorithm as proposed in Habeck (2012). This achieves an approximation of the maximum entropy Boltzmann probability density, which can be used for MCMC parameter estimation. With each iteration step, more simulated conformations are available to estimate the partition function, thus successively improving the estimate. In this sense, our method converges to the maximum entropy ensemble similar to replica averaging, but the limit of infinitely many iterations required in our approach can be systematically improved by simply increasing run time of the algorithm. In replica averaging, the maximum entropy limit is theoretically attained fr infinitely many replicas, which cannot easily be achieved by increasing the run time. However, our *repeated simulations* method described above is limited to short MD run times and is therefore insensitive to long time scales.

A remarkable feature of our algorithmic framework is that it can be modified in a straight forward manner to yield a second method, which we call the *reference ensemble* method. Here, one uses a set of given conformational ensembles, e.g. from one or several long MD trajectories. In this version, no repeated MD simulations are performed which makes our reference ensemble method similar in spirit to reweighting approaches. This means that it can only be expected to yield good results if the reference ensemble is already fairly close to the desired maximum entropy ensemble. However, in contrast to the repeated simulations version of our approach, it can be applied to long MD trajectories and therefore take conformational dynamics on long time scales into account. Since the two versions of our method have complementary strengths and weaknesses, it can be beneficial to first use the multiple simulations version to get an initial estimate for the maximum entropy coupling parameters, then run a long MD with these parameters and apply the reference ensemble version of our approach to the trajectory from this MD.

In the present exposition, we focus on atomic distance measurements and specifically on the nuclear Overhauser effect (NOE). In NOE spectroscopy, one determines the inverse sixth power of distances between hydrogen atoms from peak intensities in the spectrum. Measurement results are usually presented in terms of intervals for the respective distances between hydrogen atoms or groups thereof. Here, we assume the simplistic error model of a uniform distribution on the given NOE intervals. A generalization to other error models, following e.g. the approach of Gull and Daniell (1978) which was expanded on in Cesari et al. (2018), is left for future work.

In Section 2 we review the principle of maximum entropy and introduce the Bayesian model we use for parameter estimation. Then we introduce our adaptive Markov Chain Monte Carlo algorithm and explain its relation to the maximum entropy principle and the WHAM estimation involved. In Section 3, we proceed to illustrate qualitative features of our algorithm on the widely studied alanine dipeptide as a very simple test case. Finally, in Section 4, we demonstrate the viability of our approach in more realistic scenarios by applying it to NOE measurements on two simple proteins, namely the trp-cage, consisting of 20 amino acids, for which 168 NOE features were measured.

## 2 Methods and Theoretical Background

### 2.1 Maximum Entropy

In the present work, we always assume a preexisting thermodynamical system, described by a Boltzmann ensemble with probability density

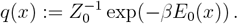

Here, *β ∈* ℝ^+^ is the inverse temperature, *E*_0_(*x*) *∈* ℝ is the energy and *x ∈* ℝ^*m*^ denotes a full description of the system under investigation, for example represented by the positions and momenta of all atoms. The energy *E*_0_(*x*) of the state *x* is given by the potential and kinetic energy of an unmodified molecular dynamics simulation and *Z*_0_ is the partition function.

The maximum entropy principle states that if an observable *y* = *f* (*x*) can only be determined in the ensemble mean 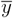, then the density function of the Boltzmann ensemble is multiplied with a term *ρ*(*f* (*x*), *θ*) := *e*^*−θf*(*x*)^, which yields the probability density

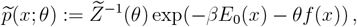

where *θ* is determined by the requirement that

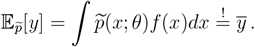

Here, we integrate over the full set of possible system states *x*. In our case of a molecular dynamics ensemble, this means an integral over the full phase space including water molecules and possible salt ions.

The maximum entropy modification is statistically least biased in the sense that the Kullback-Leibler divergence to the original ensemble is minimal and thermodynamically reasonable by maximizing entropy among all possible ensembles which are compatible with the measured quantity, see Jaynes (1957). The parameter *θ* is a coupling constant corresponding to the measured quantity. Examples include the inverse temperature *θ* = *β* corresponding to the energy *y* = *E* and the chemical potential *θ* = *µ* corresponding to the particle number *y* = *N* .

In the following, we consider *y* = (*y*^(1)^, …, *y*^(*k*)^)^*T*^ and *θ* = (*θ*^(1)^, …, *θ*^(*k*)^)^*T*^ to be vectors to enable a compact description of multiple measured features. We will write vector components as superscripts in parentheses throughout, while superscripts without parenthesis are powers as usual. In order to estimate the parameter vector *θ* from measured ensemble means 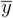, one would like to employ a Bayesian approach to properly take into account statistical uncertainty. In this context, one rewrites the modified Boltzmann density as

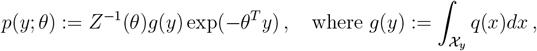

by integrating up over the classes of states 𝒳_*y*_ := *{x*|*f* (*x*) = *y}*. Then, *p*(*y*; *θ*) is used as a likelihood, and one imposes a prior distribution *π*_0_(*θ*) which expresses prior knowledge on the system. The posterior is thus given as

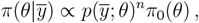

up to normalization, where *n* denotes the sample size of the measurement. In the case of typical spectroscopic measurements on biomolecules, the sample size is not an easily definable quantity, thus *n* can be understood as a parameter which expresses the relative confidence in the data versus the prior distribution. Posterior samples can be generated using an MCMC type algorithm. This approach is hindered by the fact that one requires the partition function *Z*(*θ*) to fully state the likelihood. For some system, the partition function can be calculated in closed form and a theoretical analysis of such systems is possible. However, in realistic biophysical systems, and especially in the description of large biomolecules via molecular dynamics simulations, the partition function *Z*(*θ*) typically cannot be easily calculated in a general setting and must be approximately estimated.

### 2.2 Adaptive Markov Chain Monte Carlo Method

In Bayesian statistics an estimation problem in which the normalization of the likelihood is unknown is called a *doubly intractable problem*, following e.g. Murray et al. (2006). Since *Z*(*θ*) is not available in closed form, standard sampling methods such as Metropolis-Hastings are no longer applicable. To overcome this issue, different *adaptive Markov Chain Monte Carlo* (AMCMC) algorithms have been proposed over the past two decades, see e.g. Atchadé et al. (2013); Habeck (2014); Liang et al. (2016); Park and Haran (2018). The main idea is to approximately sample from the posterior by using a proxy for *Z*(*θ*), which is updated by iteratively taking into account features sampled from the data model at different parameter values. In this article, we follow the approach put forth by Habeck (2014) which relies on sampling states *x* from the Boltzmann ensemble at each new value of *θ* by running an appropriately restrained MD simulation and thus repeatedly estimating *Z*(*θ*) by the weighted histogram analysis method (WHAM) as described in Habeck (2012) after every MD simulation. The iteration is visualized by a flow chart in Figure 1. In that Figure and in the following, subscript indices always indicate the step number of the AMCMC. In the double subscripts *y*_*mt*_, *t* indicates the AMCMC step and *m* enumerates the protein structures generated by the MD in this time step.

**Figure 1:**
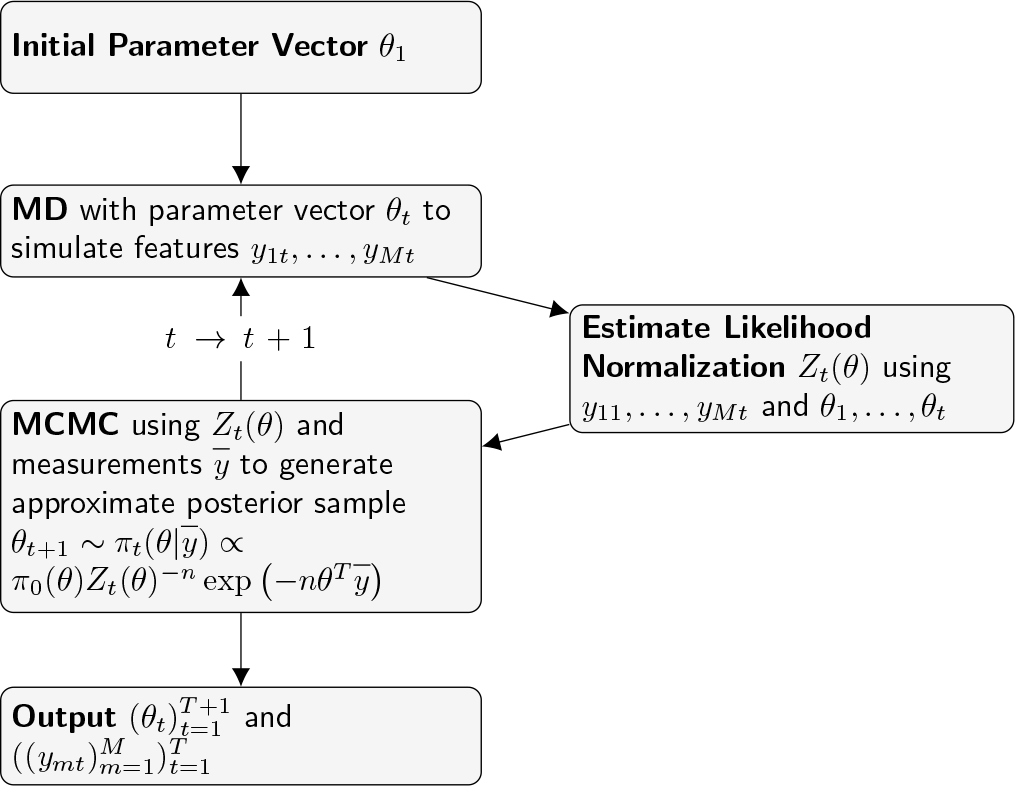
Illustration of the AMCMC algorithm proposed by Habeck (2014), which is applied and adapted in the present paper for molecular dynamics ensemble refinement.

The algorithm illustrated in Figure 1 employs repeated short MD simulations. This has the downside this it is insensitive to slow conformational changes. This limitation is especially problematic in case of negative values of parameters *θ*^(*j*)^. This is because a negative value of *θ*^(*j*)^ corresponds to a repulsive force in the MD simulation. However, since the energy term for a distance feature *y*^(*j*)^ := *r*^(*j*)^ is proportional to *θ*^(*j*)^*r*^(*j*)^, the corresponding force is independent of the distance and thus extremely long ranged. If such a repulsive force leads to a non-vanishing probability of partial unfolding of a protein, this unfolding might worsen progressively once it has started in a long simulation with fixed parameters, because stabilizing short range interactions like hydrogen bridge bonds never rejoin once they are broken. We illustrate this problem in Appendix D.

We propose a modified version of our algorithm which does not run repeated MD simulations but uses a set of fixed ensembles generated by MD runs employing different fixed parameter values *θ*^*∗k*,(*j*)^, where here and in the following, the superscript with the star enumerates the fixed ensembles. Features are then drawn from the fixed trajectories with maximum entropy weights 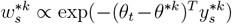 where *t* is the AMCMC time point and *s* is an index that runs over the states of trajectory *k*. The trajectories themselves are weighted with *v*^*∗k*^ *∝* exp(*−*0.5(*θ*_*t*_ *−θ*^*∗k*^)^2^*/σ*^2^) where the parameter *σ* is typically chosen close to the maximum nearest neighbor distance between the fixed parameter vectors *θ*^*∗k*^. This approach has the similar limitations as reweighting approaches. It can only yield meaningful results if at least one of the fixed ensembles used has a significant overlap with the target ensemble. However, the possibility to use multiple reference trajectories generated with non-zero parameter values *θ*^*∗k*^ makes this approach potentially more flexible than conventional reweighting approaches.

When trying to apply the AMCMC approach outlined above to NOE measurements, one encounters several complications. Firstly, NOE features are typically given in terms of feature bounds, which indicate measurement uncertainty. We use a simple approach to model a uniform probability distribution for each distance within the given range. Secondly, the peak intensity is proportional to *r*^*−*6^, such that appropriate features should be *y* = *r*^*−*6^ instead of *y* = *r*. For now, we treat these features as *y* = *r* features in the repeated simulations algorithm as is common also for methods employing flat bottom restraints. A full treatment of *y* = *r*^*−*6^ features is left for future work. For the fixed ensemble variant of our AMCMC, we restrict to reference trajectories generated at *θ*^*∗*^ = 0. Thirdly, the number of features can easily exceed 1000, in which case the AMCMC algorithm takes long to converge. We solve this problem with a divide-and-conquer algorithm by achieving convergence first on subsets of the features and then stepwise merging of these subsets. The details of all these approaches are explained in Appendix A.

In order to facilitate the interpretation of values of *θ*^(*j*)^, note that the unit of *θ*^(*j*)^ is nm^*−*1^ in the case *y* = *r*. Using the fact that *T* = 300*K* in all systems considered in this article, this can be translated to alternative widely used units as follows:

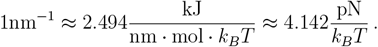

Recall that *k*_*B*_*Tθ*^(*j*)^*y*^(*j*)^ is the energy term which arises from the maximum entropy term, so it gives rise to a force 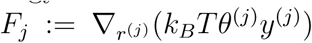. In the case *y*^(*j*)^ = *r*^(*j*)^ we get *F*_*j*_ = *k*_*B*_*Tθ*^(*j*)^ and a positive value of *θ*^(*j*)^ corresponds to an attractive force.

Accordingly, in the case *y* = *r*^*−*6^, the unit of *θ*^(*j*)^ can be written as

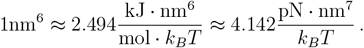

However, note that forces in this setting are *F*_*j*_ = *−*6*k*_*B*_*Tθ*^(*j*)^(*r*^(*j*)^)^*−*7^, so they incur an additional factor *−*6 from the gradient of the energy, which means that in this case a positive value of *θ*^(*j*)^ corresponds to a repulsive force. The forces arising for *y* = *r*^*−*6^ features have the same form as the attractive part of a Lennard-Jones potential, thus parameters determined in this setting can be compared to Lennard-Jones force constants for a heuristic interpretation of their size.

### 2.3 MD Parameters and Simulation Systems

All molecular dynamics simulations are done with GROMACS, using the amber99SB-ILDN force field and the TIP3P water model. The simulations use a time step of 2fs, linear center of mass motion removal every 100 steps, and energy calculation every 100 steps. We use a Verlet cut-off scheme for neighbor searching, the Particle-Mesh Ewald (PME) approximation for the Coulomb field, and the cut-off method for van der Waals interaction and apply long range dispersion corrections for energy and pressure. We use a velocity rescaling thermostat for the whole system with time constant 100fs and temperature 300K. We also use a C-rescaling isotropic barostat with time constant 1ps, compressibility 4.5 · 10^*−*5^bar^*−*1^ and a reference pressure of 1bar. Constraints are treated with the LINCS algorithm considering h-bonds.

Finally, since we use GROMACS’ distance restraint API to apply modifications to the force field for NOE features, it is important to detail the settings we use for these. We use simple individual molecule, instantaneous restraints with an energy conservative force weighting. The logging of distance restraint energies has been reduced to once every 1000 steps in keeping with logging of other energy terms.

For all other parameters, we use GROMACS default settings.

## 3 Ensemble Refinement in a Simple Test System

As a first sanity check for the two variants of the maximum entropy AMCMC we consider a simple test system comprising the alanine dipeptide in water at 300K. An alanine dipeptide molecule, depicted in Figure 2a consists of one alanine molecule, bound to an acetyl N-terminus (ACE) and a methylamide C-terminus (NME). It therefore features the Ramachandran angles *ϕ* and *ψ* exactly once and can be seen as an extremely simplified model for a single residue in a protein chain. Despite its structural simplicity, it exhibits some conformational variability which we exploit here.

**Figure 2:**
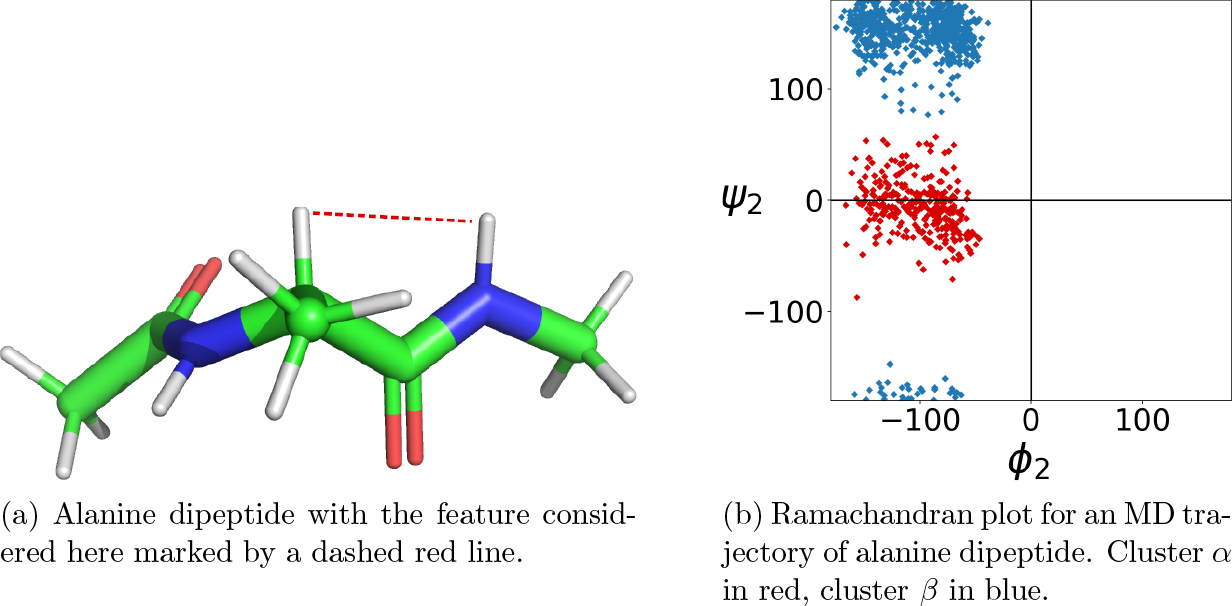
Structure and conformational distribution of alanine dipeptide.

As the single feature *y* we consider the distance *r* between the H*α* atom on the alanine and the hydrogen atom attached to the nitrogen at the methylamide C-terminus, marked by a dashed red line in Figure 2a. It is well known that alanine dipeptide has two major conformational clusters, each of which can be decomposed into two subclusters. For simplicity, we will only consider two clusters defined by the dihedral angle *ψ*. The cluster with *ψ ∈* [*−*90^*°*^, 90^*°*^] and with feature value 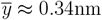 will be considered the *α* cluster because it roughly corresponds to an *α* helix fold. The other cluster with feature value 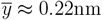 will be considered the *β* cluster because it roughly corresponds to an *β* sheet fold. The Ramachandran plot with the two clusters highlighted in different colors is displayed in Figure 2b.

The parameters used for the MD simulations are detailed in Appendix B. For an MD simulation with *θ* = 0 we get roughly a population of 35% in the *α* cluster and 65% in the *β* cluster. We perform simulations with target features 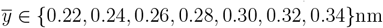 with the repeated simulation AMCMC and evaluate the resulting parameter and feature distributions in terms of kernel density estimates. The results are displayed in Figure 3a and d. In turns out that the feature distributions reproduce the two cluster structure, where different values of 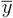 lead to different relative populations of the clusters but no unphysical structures in energetically unfavorable regions of the Ramachandran plot. A high value for 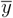 leads to a low value of *θ* and an ensemble which strongly favors the *α* cluster. Conversely, a low value for 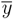 leads to a high value of *θ* and an ensemble which strongly favors the *β* cluster. For 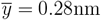 the relative weights of the clusters are close to equal. In all cases, intermediate states remain rare, so the maximum entropy potential does not enforce unphysical conformations.

**Figure 3:**
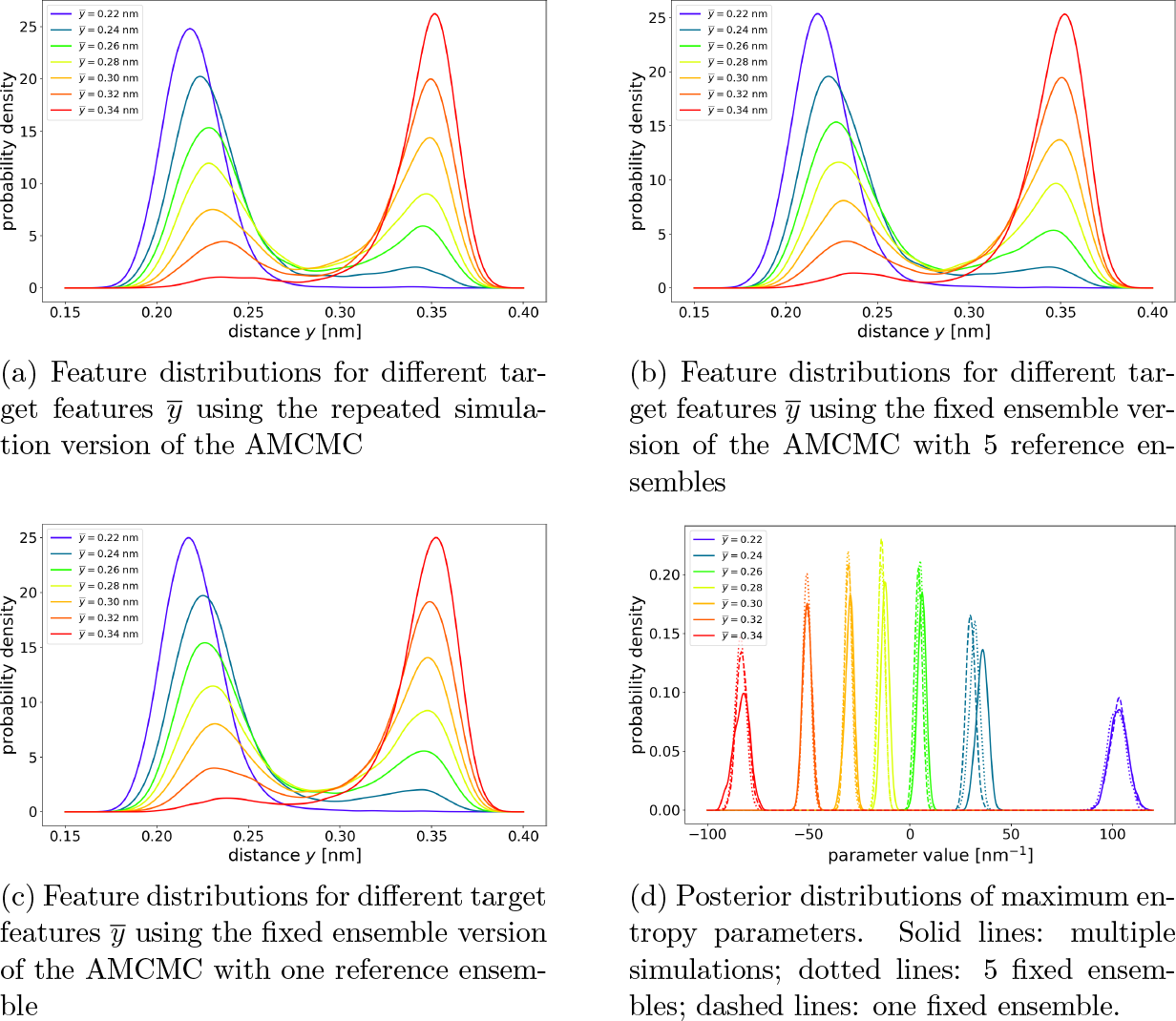
Results of three variants of the maximum entropy AMCMC for alanine dipeptide.

For comparison, we perform the same analysis with the fixed ensemble variant of our approach. We compare one variant with five reference ensembles generated with maximum entropy potentials for parameter values *θ*^*∗*^ *∈ {−*100, *−*50, 0, 50, 100*}*nm^*−*1^ with a simpler variant only using the ensemble for *θ*^*∗*^ = 0. It turns out that both variants lead to very similar posterior distributions of parameters and ensembles, compared to each other and also compared to the results of the multiple simulations version of the AMCMC. This result is not unexpected, since in this case the standard MD already generates an ensemble containing all relevant conformations. Therefore, the fixed ensemble AMCMC yields good results and does not gain much power by including ensembles for multiple values of *θ*^*∗*^.

We also apply the fixed ensemble variant of our approach using the feature *y* = *r*^*−*6^ and the target feature values 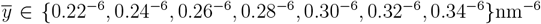 with only one fixed ensemble for *θ*^*∗*^ = 0 in Figure 4.

**Figure 4:**
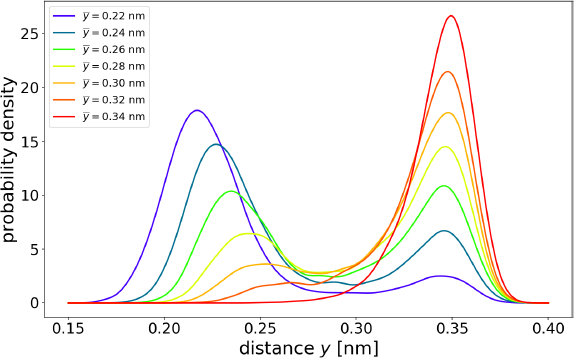
Feature distributions for different target features 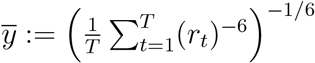 using the fixed ensemble version of the AMCMC with one reference ensemble. The cluster of structures with high distances has higher populations than for 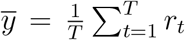 and the cluster of structures with small distances is shifted further to small distances. These two effects balance each other out to attain the target feature in the *r*^*−*6^-mean.

The cluster of conformations with larger values of *r* is weighted slightly higher than in the case *y* = *r*, especially in the case of low target distance. This is because they do not disturb the mean of features *y* = *r*^*−*6^ as much as they do the mean of features *y* = *r*. This illustrates a relative insensitivity of NOE measurement values to parts of the ensemble with large distances.

The cluster with smaller values of *r* is shifted more strongly according to the value of the target feature. For small target distances, the cluster is shifted to small distances, for larger target distances, it is shifted to larger distances. This is a consequence of the fact that short distances contribute very strongly to the mean of features *y* = *r*^*−*6^. This behavior can be expected to hold more generally, when using features *y* = *r*^*−*6^. Conformations with large distances are suppressed less than for features *y* = *r* and the distance distributions in conformational clusters at high distances are not strongly affected. In contrast, conformational clusters at small distances are not only affected in terms of the number of conformations in the cluster but also the distance distribution of the cluster itself and thereby its mean and variance.

## 4 Application to NOE Measurements

### 4.1 Trp-cage

We apply the repeated simulation version of our algorithm with features *y* = *r* to the trpcage peptide presented in Neidigh et al. (2002) available from the RCSB PDB under the id 1L2Y. The trp-cage consists of 20 amino acids comprising 304 atoms. It is widely used as a test system for small peptides, partly due to its fast folding time and highly preserved folded form, see Lindorff-Larsen et al. (2011). Its folded form has a comparatively low *α*-helix and *β*-sheet content and thus features a fairly high diversity of conformational features.

The structures given in the PDB are fits to NMR measurements and are accompanied by a set of 168 NOE features, which are presented in terms of distance ranges. In Appendix C, we explain in detail how we interpret those. The peptide has charged termini, one lysine, one arginine, and one aspartic acid, leading to a net charge of +1. To neutralize the charge and add a physiological salt concentration, we add a concentration of 0.1M/l of NaCl to the model system, which is otherwise using water as solvent and is simulated at 300K. Further parameters used for the MD simulations are detailed in Appendix B.

As is common practice, the NOE features are given in terms of intervals to indicate measurement uncertainty but no explicit error model is provided. A common way of approximately achieving feature averages lying in the given feature intervals is to use *flat bottom restraints*, which consist of adding potentials which are quadratic in the deviation from the target feature range. This means that within the given feature range, no additional force applies, while violation of the feature range lead to a force which is proportional to the distance from the target feature range and pushes the molecule in the direction of the target feature range. On the one hand, this allows for considerable systematic violations, sometimes over 0.05nm, of target feature ranges while on the other hand unnaturally restricting the ensemble by enforcing the target feature range not on the whole ensemble but softly imposing it on every single structure. Despite its drawbacks, the flat bottom ensemble serves as a useful point of comparison for our method.

In order to generate a structure ensemble from the AMCMC, we run the algorithm until the posterior distribution of the parameters has stabilized. Then we run the algorithm for an additional 1000 steps and save the final structures of every MD trajectory. From these 1000 structures we then randomly sample 100 structures. As a last step, we perform an energy minimization without restraints on all structures in vacuum to reduce small scale structure flaws like bond length and angle violations which occur in consequence of the simulation in solvent at finite temperature and affect unrestrained MD simulations equally. The resulting 100 structures are displayed in Figure 5.

**Figure 5:**
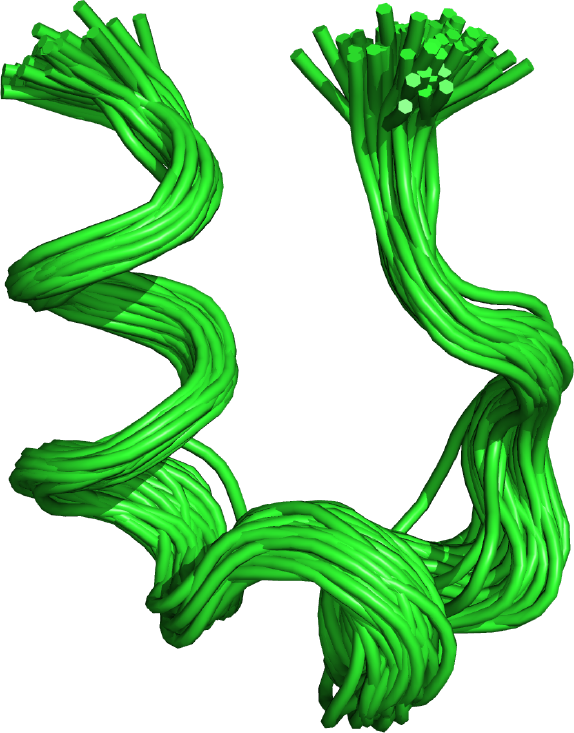
A visualization of the maximum entropy ensemble of 100 trp-cage structures. One can see that the major structural motif of trp-cage is preserved but there is fairly large variation in the structures. There are a few outliers which depart more noticably from the mean.

For the unrestrained MD, we ran one simulation for 100ns starting from a randomly chosen NMR structure and drew 100 random structures from the last 90ns of the trajectory. For the flat bottom potential, we ran 10 simulations of 100ns length starting from randomly chosen NMR structures and sampled 10 random structures from the last 90ns of each trajectory, leading again to an ensemble of size 100.

Since the termini can move fairly unrestricted, they show large variations over the data set and typically dominate the first few principal components of an all-atom PCA. To avoid this source of noise, we exclude the termini from consideration and perform PCA over all atoms of residues 2-19. The results for the first four principal components is displayed in Figure 6. In the first two principal components, one can clearly see that the flat bottom ensemble remains fairly close to the unrestrained ensemble and barely overlaps with the NMR ensemble, while the maximum entropy ensemble spans a wide range covering the NMR ensemble and part of the unrestrained MD ensemble. In principal components 3 and 4, one can see that the flat bottom potential is moved somewhat away from the unrestrained MD ensemble and in the direction of the NMR and maximum entropy ensembles. However, the flat bottom ensemble exhibits lower variance, which reflects the restriction on conformational diversity imposed by the flat bottom restraints.

**Figure 6:**
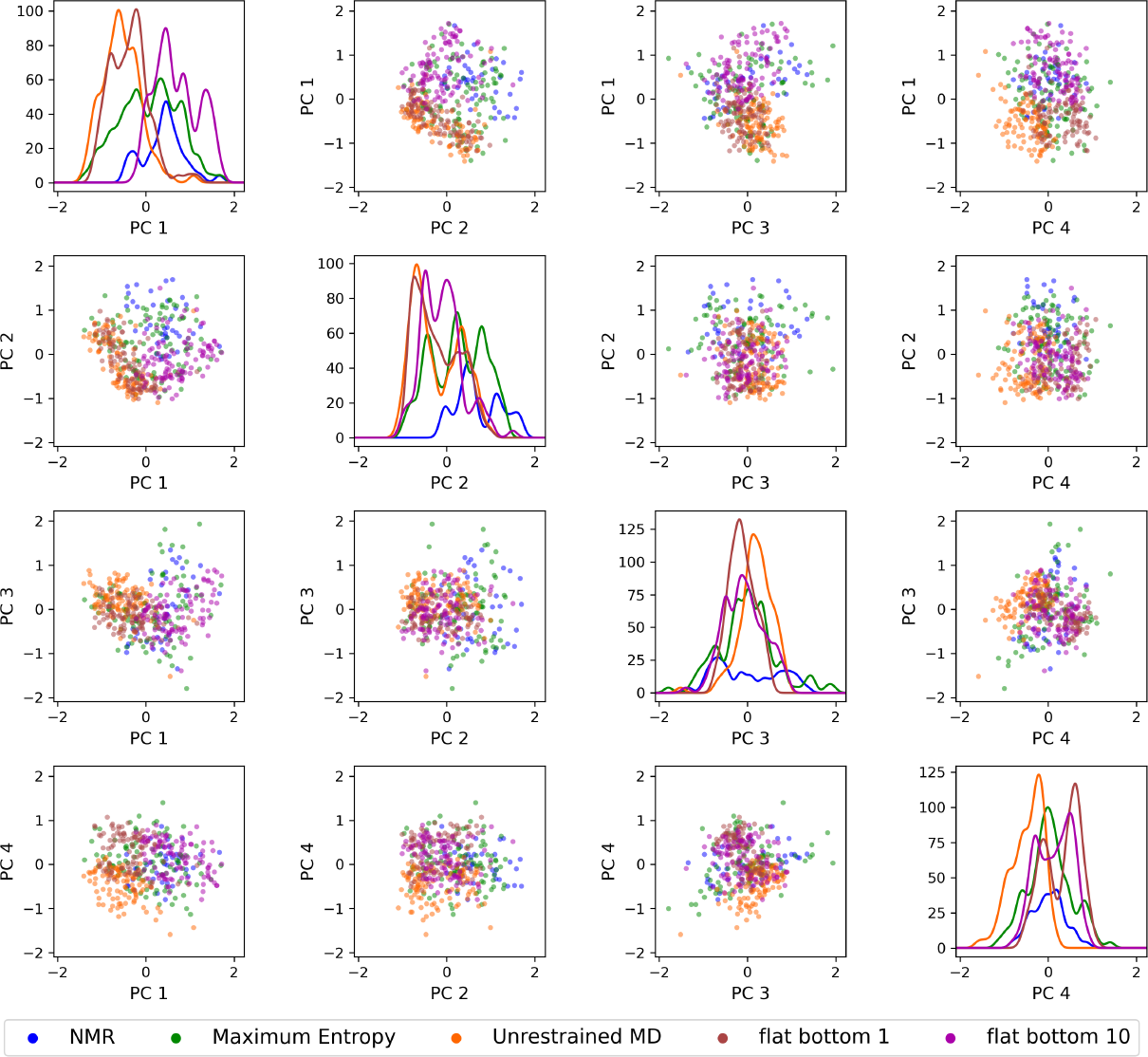
An overview of the first four principal components of the trp-cage ensembles. The PCA is performed over all atoms of residues 2-19. All four simulated ensembles consist of 100 structures. Panels on the diagonal display kernel density estimates of the individual principal component scores for the ensembles while off-diagonal panels display two-dimensional scatter plots of pairs of principal components. The axes are scaled in nanometers.

The results for the flat bottom restraints are somewhat surprising. On the one hand, it appears to remain closer to the unrestrained ensemble than the maximum entropy ensemble, which contradicts the expectation that the maximum entropy ensemble should have a minimal Kullback-Leibler divergence with respect to the unrestrained ensemble. On the other hand, the flat bottom ensemble barely overlaps with the NMR ensemble, indicating that the structures are systematically different from NMR structures. It turns out that the flat bottom potential with standard force constant is weak enough to allow for systematic NOE bound violations of around 0.05nm for several features. This explains the systematic deviation of the flat bottom ensemble from the NMR ensemble and its bias toward the unrestrained ensemble.

We have therefore performed MD simulations with flat bottom potentials with a 10 times stronger force constant. The PCA results, displayed in Figure 6, indicate that the flat bottom ensemble is shifted in the direction of the NMR ensemble. While the NOE bounds are satisfied very well on average with deviations of less than 0.01nm, the ensemble shows less overlap with the unrestrained ensemble than the maximum entropy ensemble while at the same time only overlapping with part of the NMR ensemble. This finding is closer to the expected result for the flat bottom potential, namely it indicates a higher Kullback-Leibler divergence to the unrestrained ensemble than the maximum entropy ensemble while at the same time showing less conformational diversity.

The method we propose explicitly determines a posterior distribution of maximum entropy parameter values. This allows us to identify regions in the protein where modifications of the MD force field are required to satisfy the given NOE bounds on average. We list all features which are significantly different from 0, i.e. where the absolute value of the posterior mean exceeds two times the posterior standard deviation, in Table 1.

**Table 1:** Posterior means of parameters which are significantly different from 0 for trpcage.

| feature number | atom group 1 | atom group 2 | Posterior mean $k_B T \theta^{(j)}$ [pN] |
| --- | --- | --- | --- |
| 2 | 1ASN HA | 4ILE HD(1-3) | 130.24 |
| 29 | 3TYR HE(1-2) | 7LEU HD2(1-3) | 88.93 |
| 44 | 5GLN HA | 8LYS HB(1-2) | -62.54 |
| 57 | 6TRP HA | 9ASP HB2 | -26.25 |
| 60 | 6TRP HD1 | 16ARG HB(1-2) | 63.16 |
| 62 | 6TRP HD1 | 16ARG HD(1-2) | 30.31 |
| 100 | 7LEU HB1 | 7LEU HD1(1-3) | -294.41 |
| 104 | 7LEU HD2(1-3) | 11GLY HN | 551.44 |
| 118 | 9ASP HB1 | 14SER HB2 | 41.92 |
| 141 | 14SER HN | 14SER HB1 | -296.53 |
| 142 | 14SER HN | 14SER HB2 | 36.34 |
| 157 | 17PRO HB2 | 18PRO HD1 | 21.69 |

Features 2, 29, 44 and 57 all concern distances between residues which are one helix turn apart in the *α* helix region. While the attractive forces generated by the positive values of *θ*^(2)^ and *θ*^(29)^ stabilize the terminal part of the helix, the other end of the helix is destabilized by the negative values of *θ*^(44)^ and *θ*^(57)^, which cause repulsive forces. Features 60 and 62 are very long range and measure the tightness of the folding in the middle of the peptide chain. The positive values of *θ*^(60)^ and *θ*^(62)^ indicate, that the unrestrained MD tends to allow for more leeway in the folding than the measured features would permit, which leads to an attractive force, stabilizing the main fold of the molecule. Features 104 and 118 are again fairly long range and concern the loop folding near the middle of the peptide chain. Again the positive parameter values lead to attractive forces, which indicate that the unrestrained MD allows for a less compact folding than the NOE features suggest. It is noteworthy that *θ*^(104)^ correlates strongly negatively with *θ*^(100)^ and *θ*^(118)^ correlates negatively with *θ*^(141)^ and positively with *θ*^(142)^. Thus, the high values of *θ*^(104)^ and *θ*^(118)^ are influenced by more local features. Finally, feature 157 is a local feature near the C-terminus and the positive value of *θ*^(157)^ leads to a slightly more rigid terminus shape.

The very pronounced negative values of *θ*^(100)^ and *θ*^(141)^, both associated to local features within one amino acid, appear somewhat implausible since the local conformations of amino acid are rather strongly stereochemically restricted. Therefore, one may suspect these parameter values to indicate too strict lower bounds for the features.

Of all these results, the negative values of *θ*^(44)^ and *θ*^(57)^ are most problematic, since they lead to a repulsive force in the helix conformation. Since we consider features *y* = *r*, the forces are independent of the distance so they are extremely long range. As explained in more detail in Appendix D, this leads to a partial unfolding of the peptide if one runs a longer simulation *>* 10ns applying the forces determined the posterior mean features. However, in reality, NOE features are *y* = *r*^*−*6^, which would give rise to much more short ranged forces. Therefore, we expect this problem to be mitigated in future work using *y* = *r*^*−*6^ features.

## 5 Conclusion and Outlook

We introduce a new approach to maximum entropy ensemble enhancement, which aims at explicitly determining Bayesian estimates for the maximum entropy parameters. It is thus the first maximum entropy method that not only determines parameter estimates but also quantifies the uncertainty of these estimates. We show that our method has similar technical advantages and drawbacks as other approaches with similar technical properties. Most prominently, the fact that an MD is run in every iteration step of the Monte Carlo procedure imposes stringent limits on the length of MD simulations which are feasible. This means that conformational changes which occur on long time scales may be overlooked by our method. This problem may be partly mitigated by running multiple simulations in parallel to enhance to probability of sampling rare events. We introduce a modified algorithm, which is similar to reweighting approaches, but has the advantage of being able to rely on reference trajectories generated by MD simulations at non-zero fixed values of the maximum entropy parameters. This can greatly improve the success chances of the reweighting, since it can start from ensembles which are already close to the target ensemble.

In the example of the trp-cage peptide, we show that our Monte Carlo approach can deal with the experimental data and leads to clearly interpretable results. The non-zero maximum entropy parameters are all associated with distinct conformational elements. A positive sign of the parameters indicates that the MD potential underestimates the density of parts of the folding whereas negative parameters point to a region of the molecule, where the MD force field causes a too dense folding. For the trp-cage, one pronounced effect is that the N-terminus is much more likely to bend toward the *α* helix than the unrestrained MD suggests.

At the present stage, our approach has two major shortcomings. Firstly, we use features *y* = *r*, similar to the approach employing flat bottom restraints, while in reality, NOE features are *y* = *r*^*−*6^. We plan to extend our approach to the more realistic *y* = *r*^*−*6^ setting in the near future and we expect that this will also serve to mitigate a major problem of our current approach, namely that repulsive forces can destabilize the protein on a large scale and lead to irreversible unfolding. Secondly, the maximum entropy principle is designed to deal with measured ensemble averages, which are given without a measure of uncertainty. Extensions of the maximum entropy principle to feature measurements with uncertainty are not straightforward to formulate in way that they are amenable to similar Bayesian treatment as proposed here. This problem will be investigated in future research.

## Acknowledgments

All authors gratefully acknowledge funding by the DFG CRC 1456 *Mathematics of Experiment*, project B02. M. Habeck gratefully acknowledges funding by the Carl Zeiss Foundation within the program “CZS Stiftungsprofessuren”.

## A Taylored AMCMC for NOE Measurements

### A.1 Feature Ranges

NOE measurement results are typically not provided in terms of raw spectrum data but as intervals for certain atomic distances. In fact, in most cases the features describe pairs of sets of atoms. Let 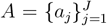 describe the positions of one set of *J* atoms in space and 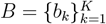 the positions of *K* other atoms. Then the corresponding NOE feature, for which bounds are given, is

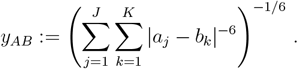

Instead of mean features 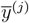, one has feature intervals [*y*^(*j*),*−*^, *y*^(*j*),+^] within which the ensemble means of the features are expected to lie. In order to accommodate this modified setting, we introduce a target feature value 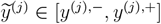 which is updated at each step of the AMCMC. If the parameter *θ*^(*j*)^ gives rise to an attractive force, 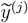 is increased, otherwise it is decreased, unless it would leave the feature interval. This means that the target features are adjusted within the feature intervals to minimize the absolute value of parameters modifying the MD and non-zero posterior parameter values only arise if the target feature reaches one of the bounds of the interval. The resulting algorithm is illustrated in Figure 7.

**Figure 7:**
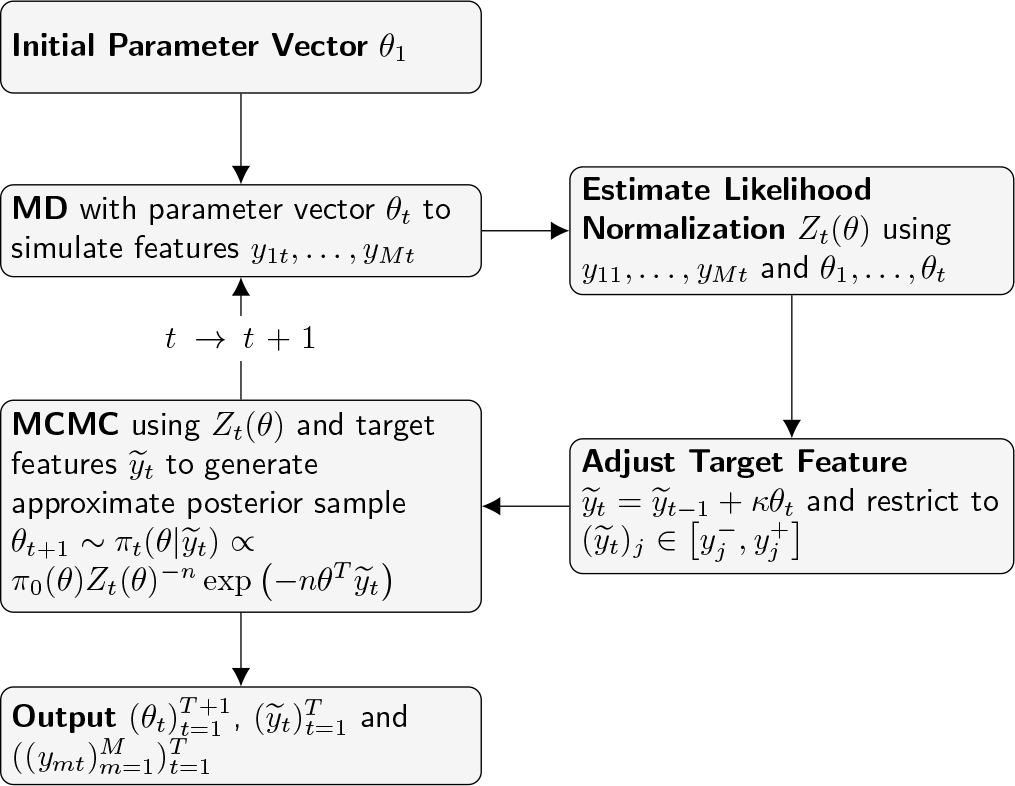
Illustration of the AMCMC algorithm taking into account feature ranges.

### A.2 Slow Convergence in Case of Many Features

If the number of features exceeds roughly 250, the AMCMC may take more than 1000 steps for posterior convergence. This is because a large set of “good” features {*y*^(*j*)^} from MD trajectories is required to estimate *Z*(*θ*) well enough to have a meaningful approximate likelihood. To deal with large feature sets, we implement a *divide and conquer* approach: the feature set is randomly partitioned into a number of smaller feature sets. The AMCMC is then run for each of these subsets until posterior parameter values stabilize. After that, the parameter and feature sets of pairs of the subsets are merged and the AMCMC is run on the merged feature sets until posterior parameter values stabilize.

This is repeated until all feature subsets are merged. Using this approach, we can achieve posterior convergence for roughly 2000 features. However, due to memory and run time requirements of the WHAM algorithm used for the estimation of *Z*(*θ*), this appears to be close to the maximum number of features for which our algorithm is viable on current hardware.

## B MD Parameters

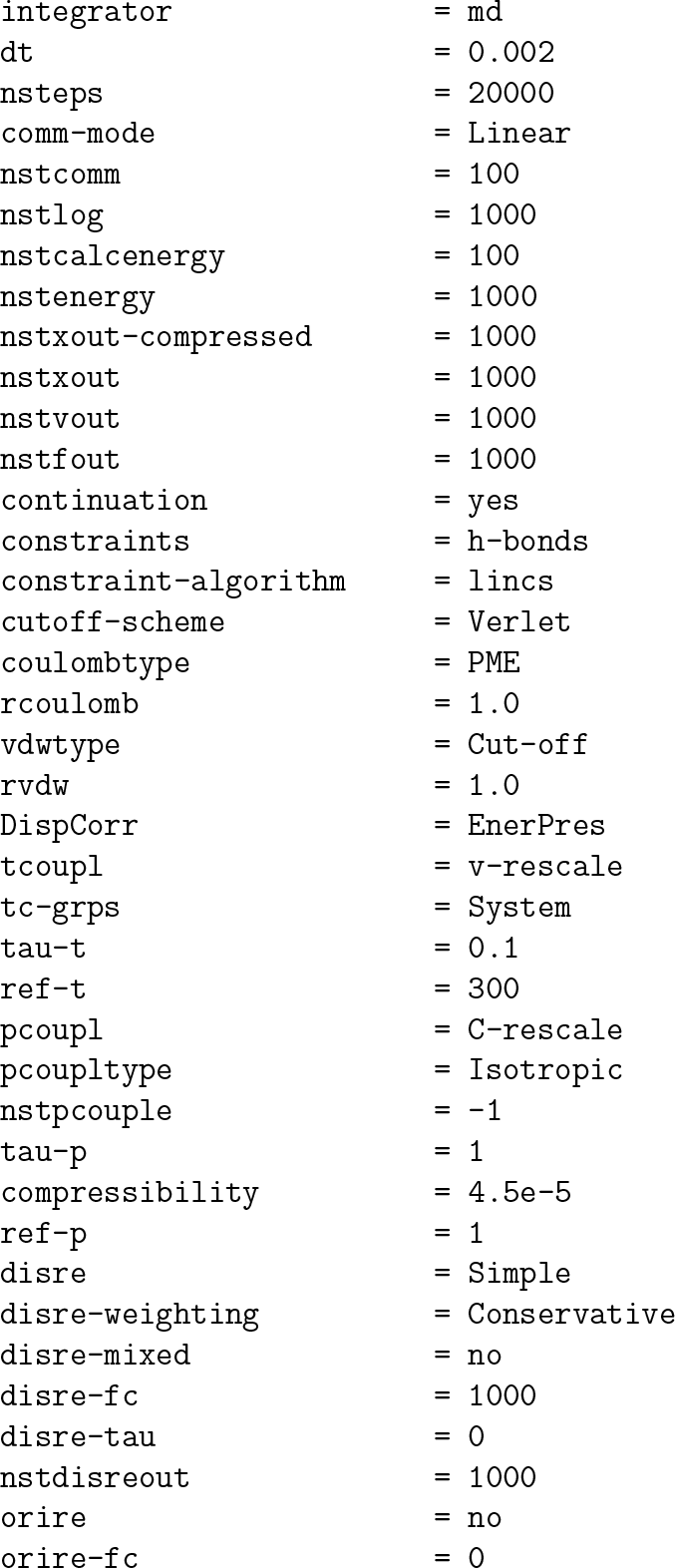
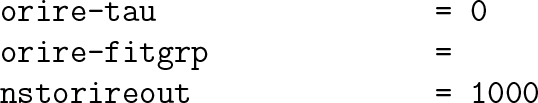

## C PThe PDB 1L2Y Data Set

The data set PDB 1L2Y has some features to be aware of. Here, we will briefly review these and explain how we deal with them.

1. The NOE feature describing the distance between the HA atom of residue 5GLN and the HN atom of residue 6TRP appears twice in identical form in the NOE data set. We have chosen to remove the duplication and work with the remaining set of 168 NOEs instead of the original list of 169.
2. The paper Neidigh et al. (2002) states that all NOEs have one of four feature bounds: 0.20-0.30nm, 0.25-0.35nm, 0.29-0.40nm, 0.33-0.50nm. However, when checking the NOE bounds, we find the following results:

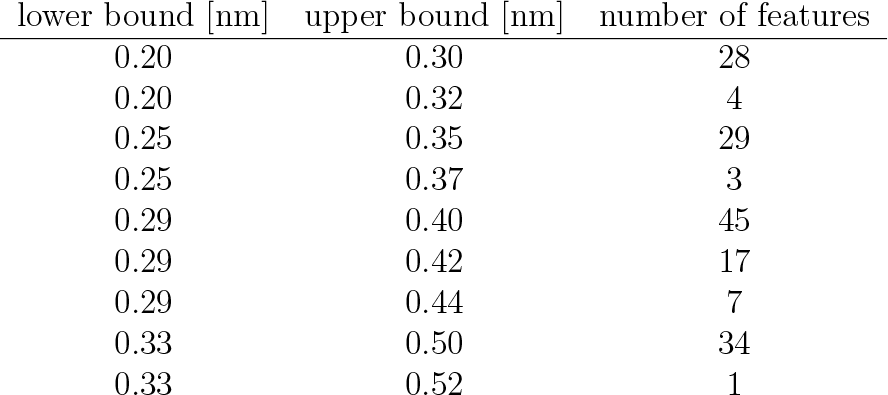

We have not found any explanation for the deviating feature bounds. It seems that all features with deviating bounds are multi-atom features, but not all multi-atom features have extended bounds. We have accepted the modified bounds as given, despite the missing justification.

3. Multi-atom-features in several cases violate lower bounds by as much as 0.05nm systematically, which is much more than stated in Neidigh et al. (2002), who claim individual bound violation to not exceed 0.011nm. This is also obvious from the PDB’s validation report. However, if one calculates multi-atom features not using the sum 1. of the inverse sixth order of distances, i.e 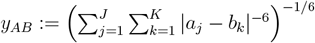, but instead using the mean, i.e.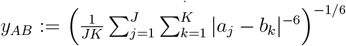, one recovers the good fit to feature bounds stated in Neidigh et al. (2002). This indicates that the authors used mean averaged multi-atom features when assessing the quality of their NOE bounds and structures. In MD simulations, it is common to instead work with sum averaged distances, 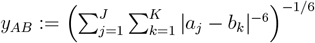, which are incompatible with the NOE bounds given by Neidigh et al. (2002). If one only applies a flat bottom potential or one only considers total bound violation over an MD simulation, this incompatibility can easily go unnoticed and it appears that this is what happened in the last 20 years. However, our AMCMC algorithm will give rise to extreme repulsive forces in order to enforce the NOE bounds of multi-atom features. In order to enable a comparison of our results to Neidigh et al. (2002), we have decided to scale all multi-atom features by 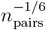 and thus apply NOE means rather than the more commonly used sum averages.

## D Repulsive Forces Destabilize Molecule Folding

In Section 4, we noted that the negative posterior mean parameters given in Table 1 can cause a local unfolding of the protein in a simulation which is much longer than the 400ps used for the MD simulations in the iteration steps of our AMCMC algorithm. In fact, simply running a long simulation of trp-cage with the posterior mean parameters, we find that the last winding of the *α* helix, closest to the middle of the peptide, unfolds within 1 *−* 5ns as shown in Figure 8. While the overall folded state of the peptide remains stable, this partial unfolding leads to significant systematic violations of multiple NOE feature bounds.

**Figure 8:**
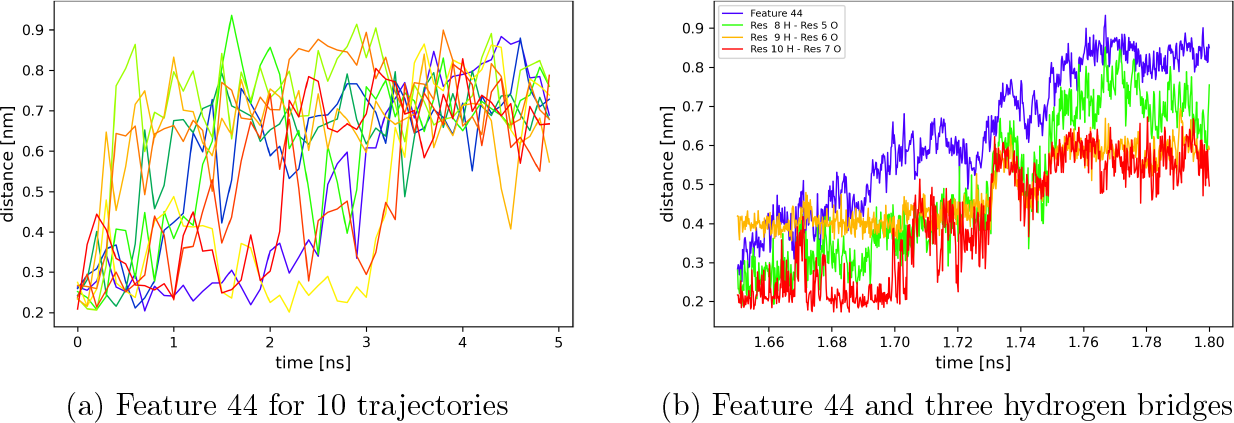
MD Trajectories of trp-cage with the posterior mean parameters applied as constant forces. Panel (a) shows the value of feature 44 (5GLN HA – 8LYS HB(1-2)) for ten trajectories starting from different conformations from the NMR ensembles. One can see that in all cases the distance increases much beyond the upper feature bound of 0.4nm and then remains high. In panel (b), one trajectory is displayed at higher time resolution and in addition to feature 44, three hydrogen bridges belonging to the most central part of the *α* helix are displayed. One can see that the growth of feature 44 precedes sudden increases in the hydrogen bridge lengths up to the point where the hydrogen bridges break.

The shape change in the helix remains irreversible over the whole duration of the simulation, since the repulsive forces are constant in time and do not depend on the distance. In the course of the partial unfolding, three hydrogen bonds in the *α* helix are broken, as illustrated in Figure 8b. One can see that the unfolding progresses quickly after the first hydrogen bond is broken.

## References

Amirkulova, D. B. and A. D. White (2019). Recent advances in maximum entropy biasing techniques for molecular dynamics. Molecular Simulation 45 (14-15), 1285–1294.

Atchadé, Y. F., N. Lartillot, and C. Robert (2013). Bayesian computation for statistical models with intractable normalizing constants. Brazilian Journal of Probability and Statistics 27 (4), 416 –436.

Barrett, R., M. Ansari, G. Ghoshal, and A. D. White (2022). Simulation-based inference with approximately correct parameters via maximum entropy. Machine Learning: Science and Technology 3 (2), 025006.

Bolhuis, P. G., Z. F. Brotzakis, and M. Vendruscolo (2021). A maximum caliber approach for continuum path ensembles. The European Physical Journal B 94 (9), 188.

Boomsma, W., J. Ferkinghoff-Borg, and K. Lindorff-Larsen (2014). Combining experiments and simulations using the maximum entropy principle. PLoS Comput Biol 10 (2), e1003406.

Bottaro, S., T. Bengtsen, and K. Lindorff-Larsen (2020). Integrating Molecular Simulation and Experimental Data: A Bayesian/Maximum Entropy Reweighting Approach. Methods Mol Biol 2112, 219–240.

Cavalli, A., C. Camilloni, and M. Vendruscolo (2013). Molecular dynamics simulations with replica-averaged structural restraints generate structural ensembles according to the maximum entropy principle. The Journal of Chemical Physics 138 (9). 094112.

Cesari, A., S. Reißer, and G. Bussi (2018). Using the maximum entropy principle to combine simulations and solution experiments. Computation 6 (1).

Crehuet, R., P. J. Buigues, X. Salvatella, and K. Lindorff-Larsen (2019). Bayesianmaximum-entropy reweighting of idp ensembles based on nmr chemical shifts. Entropy 21 (9), 898.

Gull, S. F. and G. J. Daniell (1978). Image reconstruction from incomplete and noisy data. Nature 272 (5655), 686–690.

Habeck, M. (2012). Bayesian estimation of free energies from equilibrium simulations. Phys. Rev. Lett. 109, 100601.

Habeck, M. (2014). Bayesian approach to inverse statistical mechanics. Phys. Rev. E 89, 052113.

Hummer, G. and J. Köfinger (2015). Bayesian ensemble refinement by replica simulations and reweighting. The Journal of Chemical Physics 143 (24). 243150.

Jaynes, E. T. (1957). Information theory and statistical mechanics. Phys. Rev. 106, 620–630.

Köfinger, J., L. S. Stelzl, K. Reuter, C. Allande, K. Reichel, and G. Hummer (2019). Efficient ensemble refinement by reweighting. Journal of Chemical Theory and Computation 15 (5), 3390–3401. PMID: 30939006.

Liang, F., I. H. Jin, Q. Song, and J. S. Liu (2016). An adaptive exchange algorithm for sampling from distributions with intractable normalizing constants. Journal of the American Statistical Association 111 (513), 377–393.

Lindorff-Larsen, K., S. Piana, R. O. Dror, and D. E. Shaw (2011). How fast-folding proteins fold. Science 334 (6055), 517–520.

Murray, I., Z. Ghahramani, and D. J. C. MacKay (2006). Mcmc for doubly-intractable distributions. In Proceedings of the Twenty-Second Conference on Uncertainty in Artificial Intelligence, UAI’06, Arlington, Virginia, USA, pp. 359–366. AUAI Press.

Neidigh, J. W., R. M. Fesinmeyer, and N. H. Andersen (2002). Designing a 20-residue protein. Nat Struct Biol 9 (6), 425–430.

Park, J. and M. Haran (2018). Bayesian inference in the presence of intractable normalizing functions. Journal of the American Statistical Association 113 (523), 1372–1390.

Pitera, J. W. and J. D. Chodera (2012). On the use of experimental observations to bias simulated ensembles. Journal of Chemical Theory and Computation 8 (10), 3445–3451. PMID: 26592995.

Roux, B. and J. Weare (2013). On the statistical equivalence of restrained-ensemble simulations with the maximum entropy method. J Chem Phys 138 (8), 084107.

White, A. D. and G. A. Voth (2014). Efficient and minimal method to bias molecular simulations with experimental data. Journal of Chemical Theory and Computation 10 (8), 3023–3030. PMID: 26588273.

